# Hypoxia has lasting effects on fast startle behavior of a tropical fish, (*Haemulon plumieri*)

**DOI:** 10.1101/109066

**Authors:** Mayra A. Sánchez-García, Steven J. Zottoli, Loretta M. Roberson

## Abstract

**Summary statement:** This study describes for the first time long-lasting behavioral effects of hypoxia on a tropical fish, the white grunt (*Haemulon plumieri*) from Puerto Rico.

Anthropogenic activities and climate change have resulted in an increase in hypoxia in nearshore ecosystems worldwide. The San Juan Bay Estuary System in Puerto Rico is one such ecosystem that has undergone an increase in hypoxic events over the past few years. We collected white grunts (*Haemulon plumieri*) from one of the estuary lagoons to study the effects of hypoxia on fast startle responses (fast-starts). We hypothesized that exposure to hypoxia would significantly decrease the frequency of fast-starts evoked by an abrupt sound stimulus. After an exposure to an oxygen concentration of 2.5 mg L^-1^ (40% of air saturation), there is a significant reduction in the frequency of fast-starts that is maintained for at least 24 h after the exposure. Exposure to a random sequence of oxygen levels of 5.0, 4.3 and 3.7 mg L^-1^ (80, 70, and 60% of air saturation) did not show a significant effect until one hour after exposure. We speculate that the lasting effect of hypoxia on fast-starts, thought to be involved in escape, will result in a greater susceptibility of the white grunt to predation. We have identified the Mauthner cell, known to initiate fast-starts, to allow future studies on how low oxygen levels impact a single cell and its circuit, the behavior it initiates and ultimately how changes in the behavior affect population and ecosystem levels.

## INTRODUCTION

Nearshore ecosystems that include estuaries and mangrove forests provide essential refuge and nursery habitats for many animals including fishes (Beck et al., 2001; Dennis, 1992; Laegdsgaard and Johnson, 1995; Nagelkerken et al., 2000). Approximately 50% of the world’s population now live in coastal zones (NOAA 2007; UNEP and UN-Habitat 2005). As a result, the water quality of these ecosystems are degraded by the loading of sediments, increased eutrophication resulting from sewage and animal wastes, and increased presence of pollutants, threatening marine biota and human health (e.g., Ahn et al. 2005; Diaz and Rosenberg, 2008; Diaz and Breitburg, 2009; Elison and Farnsworth, 1996; Ellis, 2006; Kennish 2002; Manciocco et al., 2014; Martinuzzi et al., 2008; Rees, 2012). One major stressor for organisms living in nearshore ecosystems is the reduction of dissolved oxygen (DO) in the water column or hypoxia. Although oxygen concentration changes naturally as a result of primary productivity, tidal flow, and seasonally variant fresh water runoff (Weis et al. 2011; Paerl et al. 1998), anthropogenic activity and climate change have increased the frequency and prevalence of hypoxic events (Diaz, 2001; Diaz and Rosenberg, 1995, 2008; Diaz and Breitburg, 2009). Tropical waters are particularly susceptible to hypoxic conditions as a result of high water temperatures that accelerate organic decomposition and deplete oxygen content (Chapman and McKenzie, 2009). Depending on the persistence of an hypoxic event, the survival of aquatic animals can be compromised, with fishes being one of the most threatened organisms (Shimps et al. 2005; Diaz and Rosenberg 1995).

Hypoxia has been shown to impact behavior in a variety of fish species (Lefrançois and Domenici, 2006; Lefrançois et al., 2005; Stierhoff et al., 2009; Wannamaker and Rice, 2000). Hypoxia results in reduced responsiveness and a change in sidedness of startle responses in European sea bass (*Dicentrarchus labrax,* Lefrançois and Domenici, 2006) and golden grey mullet (*Liza aurata,* Lefrançois et al., 2005). Since startle responses are thought to be important in escape from predation, hypoxia may have adverse effects on population size, leading to an overall destabilization of an ecosystem (Breitberg, 2002; Domenici, et al., 2007; Kennish 2002). We wondered whether hypoxia affects fast startle responses (fast-starts, Domenici and Blake, 1997; Eaton et al., 2001) of a tropical fish, the white grunt (*Haemulon plumieri*), and whether the effects continue once fish are returned to normoxia or saturated oxygen conditions. We chose to study the white grunt, because it is an abundant tropical species (Courtenay, 1961; Darcy, 1983) and it is an important ecological, commercial and recreational fish throughout the Caribbean (De Silva and Murphy, 2001). Additionally, this fish is used as a bio-indicator for water quality by the Mesoamerican Barrier Reef System (MBRS) Synoptic Monitoring Program (Alpuche-Gual and Gold-Bouchot, 2008).

White grunts were collected from Condado Lagoon in the San Juan Bay Estuary (SJBE), the largest estuary in Puerto Rico with a legacy of uncontrolled urban expansion and pollution that has threatened the health of this ecosystem for decades (Fig. 1A, B; Kennedy et al., 1996; Webb and Goméz, 1998). The SJBE, located within the metropolitan area, was designated by the U.S. Environmental Protection Agency National Estuary Program (NEP) as “an estuary of national importance” due to its ecological and commercial importance (Otero and Meléndez, 2011) and is the only tropical estuary within the NEP. Our results indicate that hypoxia lowers the frequency of fast-starts in white grunts and more importantly, continues to disrupt fast-starts beyond the hypoxic treatment. We have identified the Mauthner cell (M-cell) of the white grunt as a first step in determining the neuronal mechanisms that might underlie the effects of hypoxia on startle responses. We discuss the implications of changes in this behavior on population and ecosystem structure.

**Figure 1.**
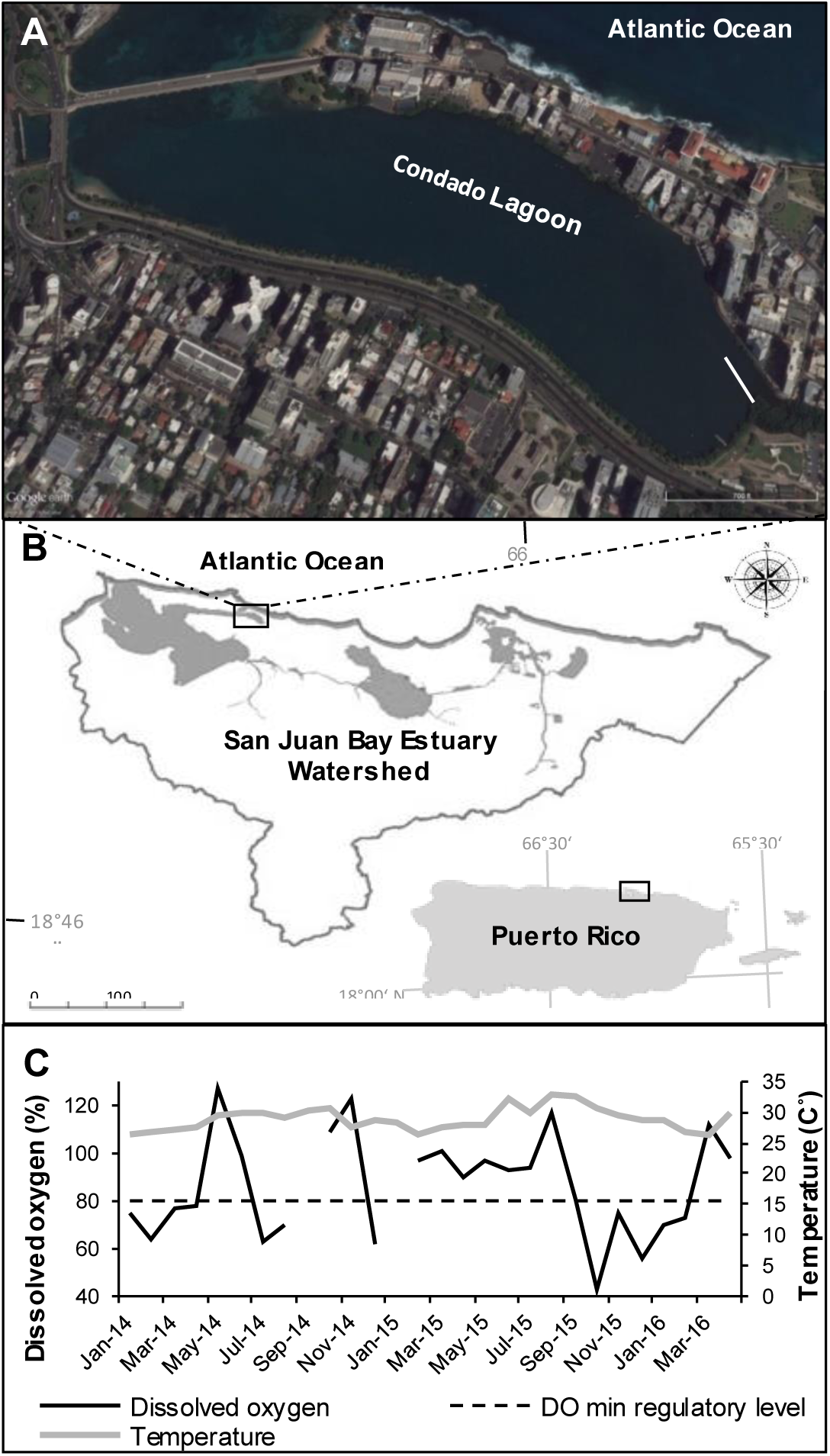
Sample collection site. **A) Condado lagoon, Puerto Rico.** The lagoon is connected to the Atlantic Ocean to the north and to the San Juan Bay by way of the San Anton channel to the west. Collection of fishes was done along a bridge that extends 100m from the eastern shore (white line) (Image: Google maps). **B) San Juan Bay Estuary.** The black square indicates the location of the Condado lagoon (Image: PRCEN). **C) Dissolved oxygen and temperatures in Condado lagoon 2014-2016.** The graph shows percent dissolved oxygen (black line) and water temperature (⁰C, gray line) during the collection period of the samples. The dashed line indicates the minimum oxygen level for healthy waterways (EPA).

## MATERIALS AND METHODS

### Collection site, fish collection, and maintenance

Specimens were collected in the Condado lagoon of the San Juan Bay Estuary from January 2014 through April 2016 (collection permits O-VS-PVS15-SJ-00595-16042013, R-VS-PVS15-SJ-00409-290814, R-VS-PV15-SJ-00482-02092015). The Condado lagoon was chosen because it is a nursery for a number of fishes including the white grunt and is subject to periodic pollutant effluence and changes in DO that result in hypoxic zones in the lagoon. Fish were collected from a pier that extends approximately 100 m from the shore on the eastern side of the lagoon (Fig. 1A). The depth of the collection site is approximately 1 m with average water temperatures in the range of 27-31˚C, and salinity between 32-40 ppt. DO ranged from 66-106 % air saturation (4.2-6.9 mg L^-1^) during the collection period (Fig. 1C).

The white grunt, *Haemulon plumieri* (9.5 + 1.4 cm, mean total length + standard deviation; range, 6-13 cm total length), were caught by cast net. Fish were transported in insulated buckets with constantly aerated water to the laboratory. Upon arrival, fish were transferred to 50 gallon holding tanks. Fish 6-9 cm in total length were housed with no more than four fish per tank, while fish 10-13 cm in total length were housed with no more than three fish per tank. The temperature and salinity of the holding tank sea water (Instant Ocean, Spectrum Brands, Inc.) was maintained within the ranges of the sea water at the collection site. Fish were exposed to an alternating 12 h light/12 h dark cycle. Water quality (i.e., salinity, pH, and temperature) was monitored daily utilizing standard methods, and nitrite and nitrate levels were measured weekly. Fish were fed three times a week with raw squid or freeze-dried shrimp (Omega one®). Any food not consumed was removed from the holding tank after two hours. Fish were observed for 3-5 days prior to experimentation to ensure they were free of infections and that they ate regularly. Fish were held for a minimum of three days prior to experiments. Before each experimental treatment, fish were deprived of food for 24 h (IACUC protocol # 00819-08-16-2013 and #01006-01-09-2015)

### Experimental set-up and image analysis

A circular plexiglass test tank (27 cm inside diameter x 19.4 cm depth) was placed on a wooden support frame on top of 15 cm speaker (TANNOY, MUSIC Group Commercial SC Ltd, Canada). The tank was filled with salt water to a depth of 10 cm (6 L). The temperature in the chamber was maintained between 27-31 °C to match the temperature at the collection site. Normoxic oxygen levels (100% DO = 6.4 mg L^-1^) were maintained by bubbling air into the water and nitrogen gas was bubbled in the water to establish hypoxic conditions. Continuous measurements of DO were made inside the test chamber with a ProODO probe (YSI, Inc.) and pH with a pH/CO2 controller (TUNZE^®^ 7074/2) during the experimental procedure and adjustments were made as needed to keep dissolved oxygen levels constant. During experiments, the pH and temperature remained constant. The outside of the test tank was covered with an opaque film and dark fabric was draped over the entire setup to eliminate visual stimuli of the fish by experimenters. The stimulus consisted of one cycle of a 100Hz signal produced by a digital waveform generator (LDB, RAG 101) in combination with an audio power amplifier (Radio Shack MPA-50, Tandy Corporation, Fort Worth, TX). A high-speed camera (The MotionXtra^®^ HG-XR Imaging System, DEL Imaging System, U.S.A) positioned above the test chamber was used to record the response of the fish at 1,000 frames per second and two hundred and fifty milliseconds of data (i.e., 250 frames) for each trial were saved for analysis. An LED light on the side of the tank provided a marker for the onset of the stimulus (Fig. 2A).

**Figure 2.**
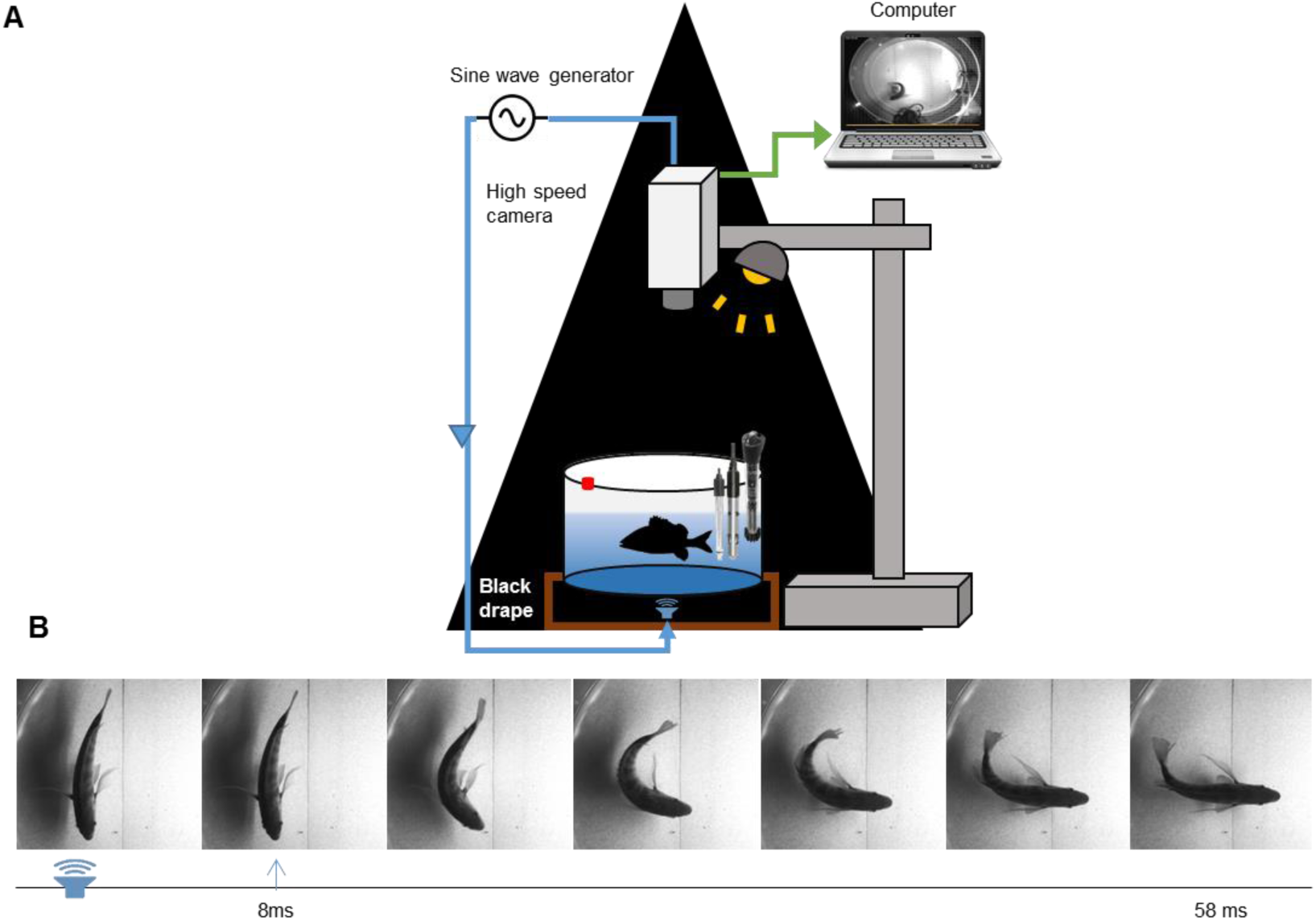
Schematic of the behavioral test arrangement. **A) Test tank set-up** (Not drawn to scale). A white grunt was placed in a test tank and after acclimation was stimulated with an abrupt sound stimulus consisting of a single cycle of 100Hz sine wave. The activation of the sound simultaneously triggered a high speed camera (1000 fps) and an LED (red square on tank). **B) A sequence of images of a fast start response (C-start).** Initial image denotes the onset (sound icon) of the stimulus followed by the first movement of the head 8 ms later (arrow). Subsequent images are spaced at 10 ms intervals.

Two variables were calculated from the imaging data: 1) frequency of fast-starts, and 2) latency of the response, as the time interval from the stimulus onset to the first movement of the head (only latencies less than 50 ms were considered fast-start responses) (Fig. 2B). We did not compare directionality of the responses between normoxic and hypoxic conditions since it is difficult to determine the directionality of the stimuli. That is, the tank sits on the speaker and thus the stimulus is distributed over the entire base of the tank.

Image analyses of the two variables were performed independently by two individuals. For frequency of fast-starts, the two individuals agreed 98% of the time. For fast-start latency, agreement occurred 76% of the time. However, differences were no more than two frames (2ms). In instances where there was disagreement among the individuals, the final value used was chosen by the most experienced recorder (i.e., M.S.G.).

The tank location of a fish prior to stimulation was recorded during each trial to ensure that position did not influence the frequency of response. The preferred positions were along the edge of the tank and the position did not affect whether fast-starts were elicited or not.

### Startle response protocols

Two separate experimental protocols were used to study the effects of low dissolved oxygen on the white grunt startle response: a single hypoxic protocol and a multiple hypoxic protocol. Both protocols consisted of three principal treatments: 1) baseline normoxia (6.4 mg L^-1^; 100% air saturation), 2) either single or multiple hypoxic conditions, and 3) a reversal from hypoxia back to normoxia. For both protocols, a single fish was placed in the test chamber under normoxic conditions and left to acclimate for a period of 30 min prior to testing. In this baseline normoxia treatment, all fish performed at least one startle response to the right and another to the left. For the single hypoxic protocol only fish that responded 80% of the normoxic trials were used in following treatments and for the multiple hypoxic treatment only those that responded 67% of the normoxic trials.

The lowest non-lethal oxygen level that caused loss of equilibrium (3 fish total) was 1.88 mg L^-1^ (30% air saturation). In comparison, equilibrium remained normal when fish were exposed to 2.5 mg L^-1^ (40% air saturation) oxygen. As a result, we selected oxygen levels of 2.5 mg L^-1^ or greater for all hypoxic treatments.

#### Single hypoxic protocol

A single exposure to an oxygen level at 2.5 mg L^-1^ (40% of air saturation) was performed to assess the effects on the frequency and latency of startle responses. Twenty-seven fish were collected; 5 were not used because they did not respond 80% of the time in the initial normoxic condition and 3 were not used because they were not tested 24 h after the hypoxic treatment. The sample size for the control group therefore was ten and the experimental group was nine. Following acclimation in normoxic conditions (6.4 mg L^-1^), experimental fish were stimulated for six consecutive trials with 3-4 min inter-trial intervals. Nitrogen was then bubbled over 15 minutes to bring the oxygen level down to 2.5 mg L^-1^ (40% DO). Each fish was then acclimated at 2.5 mg L^-1^ for 10 min before stimulation. After six trials (approximately 18 min), air was bubbled for 15 min to bring the DO concentration back up to 100% saturation where it was held prior to testing. Fish spent 18 min in oxygen levels of 2.5 mg L^-1^ (40% DO) and 30 min in partial hypoxic conditions (i.e., shifts between oxygen treatments). After the normoxia-hypoxia-normoxia sequence, fish were returned to their home tank and 24 h later were brought back to the test tank, acclimated for 30 min under normoxic conditions and tested again. Fish were then returned to the holding tank and observed over 2-3 days to ensure treatment did not adversely affect fish equilibrium and/or their ability to feed, and then they were returned to Condado lagoon. The protocol is graphically represented in Fig.3 A.

**Figure 3.**
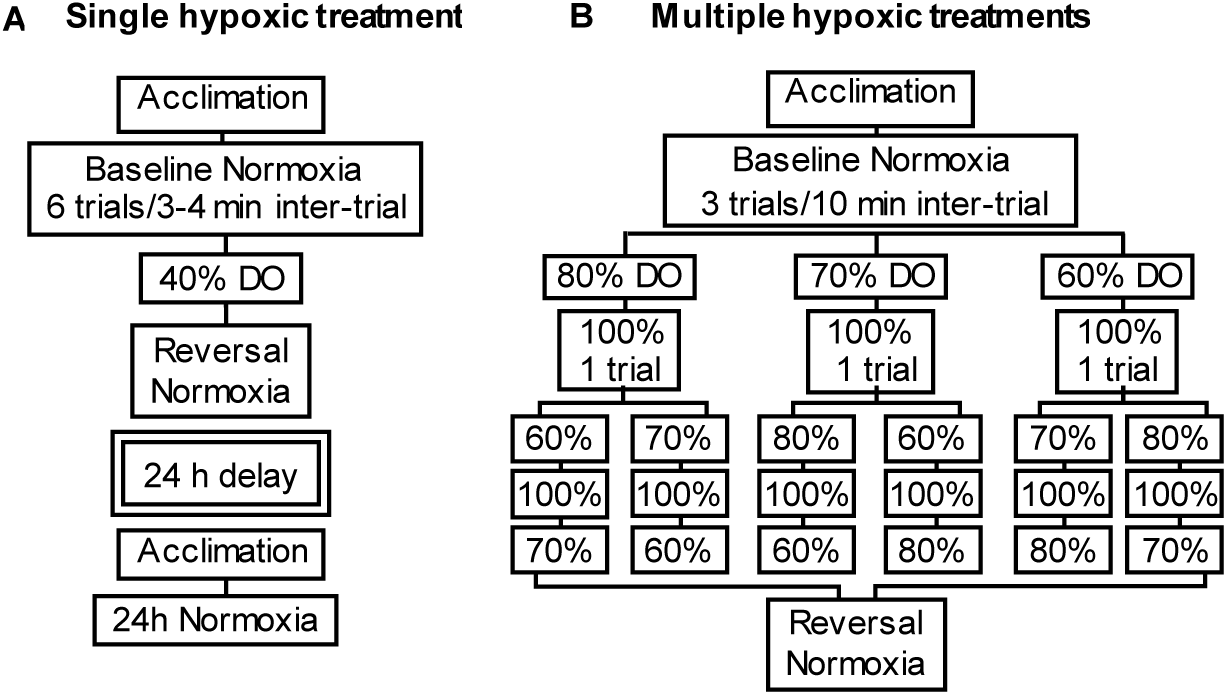
Flowcharts of single and multiple hypoxia protocols. Both protocols consisted of three principal treatments: 1) baseline normoxia (100% DO), 2) hypoxic conditions (either single or multiple levels), and 3) a reversal from hypoxia back to normoxia. **A) Single hypoxic protocol.** Each fish was placed in the test tank to acclimate in normoxic conditions and then startle responses were measured in six consecutive trials with 3-4 min inter-trial intervals. The fish were then exposed to an oxygen level of 2.5 mg L^-1^ (40% DO) and tested again. The water in the test tank was then brought back to normoxia (reversal normoxia) and fish response was tested. The fish were then returned to their holding tank and tested 24 h later (24 h normoxia). **B) Multiple hypoxic protocol.** Each fish was acclimated to normoxia and then stimulated three times with 10 min inter-trial intervals. Each fish was placed in one out of six randomly chosen oxygen concentration sequences (e.g., 5.0, 4.3 and 3.7 mg L^-1^, 4.3, 5.0, 4.3 and 3.7 mg L^-1^ etc.) and 3.7 mg L^-1^). Fish were tested at each DO and in between treatments the water was brought to normoxic levels. After exposure to three hypoxic treatments, the water in the test tank was brought back to normoxia and fish were tested again.

Control fish were subjected to the same intervals and treatment times as experimental fish but were maintained under normoxic conditions for all trials. The same aeration sequence was used except that air was bubbled instead of nitrogen. Control fish were not tested 24 h after the last normoxic treatment.

#### Multiple hypoxic protocol

A random sequence of exposures to oxygen levels of 5.0, 4.3 and 3.7 mg L^-1^, (80, 70, and 60% of air saturation) was used to assess the effects of less severe hypoxia on the frequency and latency of startle responses. Of the twenty fish that were collected one died and one were not used because they did not respond 67% of the time in the initial normoxic treatment. As a result, the sample size for the control group was nine and the experimental group was nine. After acclimation and testing under normoxic conditions, experimental fish were subjected to a randomized order of hypoxic treatments (Fig. 3B).

Each fish was placed in one out of six randomly chosen oxygen concentration sequences (e.g., 5.0, 4.3 and 3.7 mg L^-1^, 4.3, 5.0, and 3.7 mg L^-1^ etc.). For each oxygen level tested, the DO was progressively lowered at a constant rate over a 15 min period and maintained at a plateau for 10 min before testing the fish three times with a 10 min inter-trial interval. To ensure that the responsiveness of the fish was not lost after each hypoxic treatment, the DO was progressively raised back to oxygen levels of 100% air saturation (normoxia) at a constant rate over a 15 min period and maintained at a plateau for 10 min before stimulation. If a fish responded in one of two trials (all fish met this criterion), then the DO was lowered over 15 min to the next hypoxic treatment. The total time the fish spent in the experimental chamber was 5.6 h.

Control fish were subjected to the same intervals and treatment times as experimental fish but maintained under normoxic conditions for all trials as in the single hypoxic treatment. The same aeration sequence was used except that air was bubbled instead of nitrogen. The final hypoxic treatment for experimental and control fish was then used to calculate the average response to multiple hypoxic treatments (see Statistics section).

### Histological Techniques

Two white grunts were used for morphological characterization of M-cells. Fish were anesthetized in 0.03% ethyl-m-aminobenzoate (Sigma, St. Louis, MO) until respiration ceased. The heart was exposed, a cannula was placed through the ventricle into the bulbous arteriosus and secured by looping and tying suture thread around the junction. Fixative (4% paraformaldehyde in phosphate buffer, pH 7.4) was then perfused through the circulatory system. The brains were removed and placed in fresh fixative overnight.

The brains were dehydrated, cleared, embedded in paraffin and sectioned in the transverse plane at 15 µm. Sections were stained with Morse’s modification of Bodian’s silver technique (see Zottoli et al., 2011), dehydrated and cover slipped.

### Electrophysiological techniques

Five white grunts were used for electrophysiological characterization of M-cells. Fish were initially anesthetized in 0.03% ethyl-m-aminobenzoate (Sigma) until respiration ceased. They were then placed in a holding chamber and secured between tapered stainless steel rods whose tips were coated with topical anesthetic (20% benzocaine in a water soluble glycol base; Ultra-Care; Ultradent Products Inc). In the holding chamber, aerated sea water containing 0.012% of anesthetic was passed through the mouth and over the gills. The skin over the skull was then coated with local anesthetic (20% benzocaine in a water-soluble glycol base; Ultra-Care). After 10 min, the skull was removed and the hindbrain exposed. Care was taken to avoid contact of the local anesthetic with the brain and spinal cord. Two hundred micrograms of pancuronium bromide (MP Biomedicals, LLC) was injected into the trunk musculature at the mid-body level about a 1-2cm ventral to the dorsal fin. Once all operations had been performed and all exposed surfaces had been coated with local anesthetic, the fish were taken off of general anesthesia for physiological recordings. Local anesthetic was reapplied to exposed tissues during the experiment at 20 min intervals.

The dissection to expose the surface of the medulla oblongata is similar to that described for the sea robin (Zottoli et al., 2011). The hindbrain was exposed from the optic tecta to the rostral spinal cord. To expose the fourth ventricle and the surface of the medulla oblongata, a portion of the cerebellum was removed and the remainder was displaced rostrally and held in place with Kimwipes™ (Kimberly-Clark Worldwide Inc., Canada). The surface of the medulla oblongata was completely exposed by separating the overlying tissue at the midline and gently displacing each half laterally. In most preparations, the M-axons were visible crossing the midline and extending laterally toward their cell of origin. The M-cell somata cannot be seen because they are approximately 200-250 µm below the surface of the medulla oblongata. The spinal cord was exposed a few centimeters rostral to the caudal peduncle and bipolar stainless-steel stimulating electrodes were placed on vertebrae over the cord to antidromically activate the M-cells. The white grunt M-cell is located approximately 300 µm lateral to the midline and at a rostro-caudal level that is approximately centered on the cerebellum. A glass microelectrode (3 M KCl, 3 MΩ) was lowered in steps into the brain to a maximum depth of 350 µm while searching for the presence of a short-latency, antidromically-evoked extracellular negative field potential. Subsequent penetrations were spaced about 50-100 µm apart in a grid-like fashion to find the maximum field potential. A field potential of 10 mV or greater was the criterion used to identify the presumed axon cap (Furshpan and Furukawa, 1962).

### Statistical analyses

For the single hypoxic protocol, frequency of response and latency for each fish were calculated as an average of the values for all trials in a treatment (i.e., baseline normoxia, 40% DO, etc.). We then used these values to calculate an average value for all fish within each treatment. Statistical comparisons of averages were made between baseline normoxia and subsequent treatments. For the multiple hypoxic protocol, frequency of response and latency for each fish were calculated as an average of the values for all trials in the baseline normoxia, the last hypoxia treatment, and the reversal normoxia treatment. We then used these to calculate an average value for all fish within each treatment and calculated the standard error of the mean with an adjustment for propagation error. Statistical comparisons of averages were made between baseline normoxia, the last hypoxic treatment and reversal normoxia. The frequency of response and latency for each group in the protocol was analyzed with a repeated measure one-way ANOVA with a Bonferroni *post hoc* analysis for data with a Gaussian distribution. For non-normal data, a Friedman one-way ANOVA was performed with a Dunn’s *post hoc* analysis. The significance level was set at 0.05. Prism 6 software (GraphPad Software, Inc. Version 6) was used for statistical analysis and graph generation.

## RESULTS

### Single hypoxic protocol

#### Frequency and latency of response

Frequency of response of the control group showed no significant difference between the three normoxic treatments (baseline normoxia, hypoxia control, and reversal normoxia) (RM one-way ANOVA, F_(2, 18)_ = 2.76, *p* = .0902, Fig. 4A1). Latency of response of control group did not show significant difference among the three normoxic treatments as well (RM one-way ANOVA, F_(2, 18)_ = .1695, *p* = .845, Fig. 4A2).

**Figure 4.**
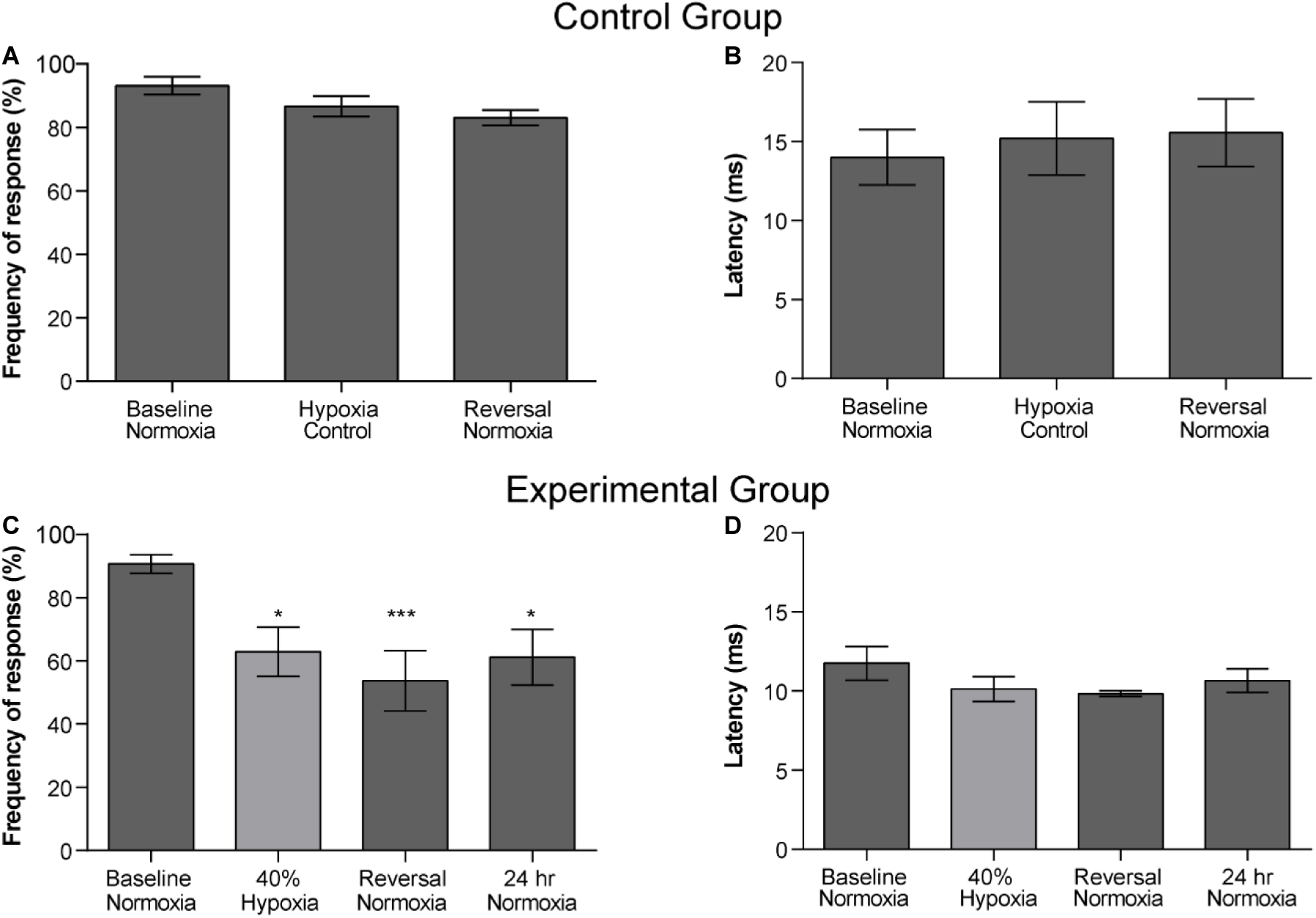
Frequency and latency of startle responses for the single hypoxia protocol. A, B) Frequencies and latencies for control fish kept in normoxic conditions throughout the protocol. A) Comparison of the frequency of response (n = 10) between the baseline normoxia and subsequent exposure to normoxia utilizing the control protocol time sequence. No significant difference was observed among treatments (*p* = 0.0902). B) Comparison of the latency of response (n = 10) between the baseline normoxia and subsequent control exposure to normoxia utilizing the control protocol time sequence. No significant difference was observed among treatments (*p* = 0.8454). C) Comparison of the frequency of response in experimental fish (n=8) between the baseline normoxia and the subsequent conditions of the single hypoxic protocol. There was a significant reduction in the frequency of response for the hypoxic treatment and normoxia reversal treatments (*p* = 0.0005). D) Comparison of the latency of response (n=8) between the baseline normoxia and the subsequent conditions of the single hypoxic protocol. No significant difference was observed among treatments (*p*= 0.2629).

Frequency of response in the experimental group was significantly reduced by exposure to 2.5 mg L^-1^ of oxygen (40% DO). Friedman’s test indicated differences between normoxia and other conditions (Friedman’s *X*^*2*^ = 17.74 df =4, n= 9, *p* = .0005, Fig. 4 B1). Post hoc comparison using Dunn’s test indicated that frequency of response decreases significantly for all treatments when compared to baseline normoxia (40% hypoxic treatment (t_B-40%_, *p* = .0244), after the reversal normoxic treatment (reversal, tB-R *P*=.0004) and 24 h later in the 24 h normoxic treatment (t_B-24h norm_, *p* = 0.0244). The latency of the response was not affected by hypoxia. No significant difference was observed among baseline normoxia, 40% hypoxia, reversal normoxia, and 24 h normoxic treatment (Friedman’s *X*^*2*^ = 3.986 df =4, n= 8, *p* = 0.2629, Fig. 4B2).

### Multiple hypoxic treatments

#### Frequency and latency of response

Frequency of response for the control group showed no significant difference between the three normoxic treatments (baseline normoxia, hypoxia control, and reversal normoxia) (Friedman test X^2^ = 4.667, df = 3 *p* = 0.222) (Fig. 5A1). Latency of response showed no significant difference as well (Friedman test X^2^ = 1.556, df = 3 *p* = 0.569, Fig. 5A2).

**Figure 5.**
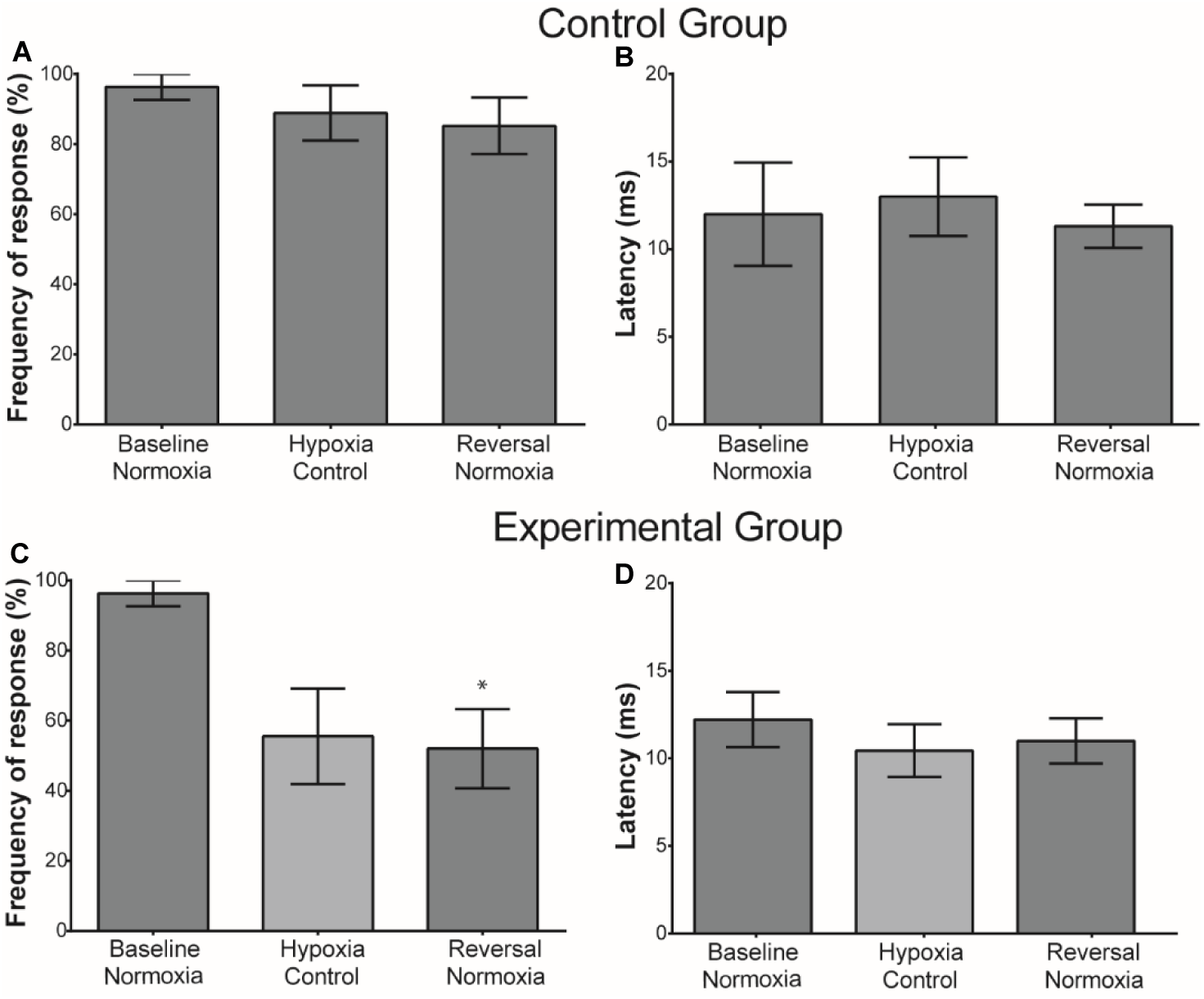
Frequency and latency of startle responses for the multiple hypoxia protocol. A1, A2)Frequencies and latencies for control fish kept in normoxic conditions throughout the multiple control protocol. A1) Frequency of response (n = 9) between the baseline normoxia and subsequent exposure to normoxia utilizing the multiple hypoxic protocol time sequence. There was no significant difference among treatments (*p* = 0.222). A2) Comparison of the latency of response (n = 9) between the baseline normoxia and subsequent exposure to normoxia utilizing the multiple hypoxic protocol time sequence. No significant difference was observed among treatments (*p* = 0.5690). B1) Comparison of the frequency of response in experimental fish (n=8) between the baseline normoxia and the last hypoxic treatment and the subsequent normoxia treatment in the multiple hypoxia protocol. ANOVA shows a significant difference among baseline and the treatments (P = 0.0057). Dunn’s post-hoc analysis shows that there was no significant reduction in the frequency of response for the last hypoxia treatment (*P* = 0.1018), however significant difference was observed on reversal normoxia treatments (*P* = 0.0401). B2) Comparison of the latency of response (n= 6) between the baseline normoxia and the subsequent conditions of the multiple hypoxic protocol. No significant difference was observed among treatments (*P* = 0.3017).

Frequency of response of the experimental group was significantly reduced when exposed to multiple hypoxia treatments (Friedman’s X^2^ = 9.923, df = 3 *p* = 0.0057, Fig. 5B1). Dunn’s Post hoc test showed no significant difference between baseline normoxia and the last hypoxic treatments (*p* = 0.1018), but did show a significant difference between the baseline normoxia and reversal normoxic treatments (*p* = 0.0401).

The latency of the response was not affected by hypoxia in those fish that responded to the stimulus. No significant difference was observed among the treatments (Friedman’s *X*^*2*^ = 2.696 df =3, n= 6, *p* = 0.3017, Fig. 5B2).

### Latency distributions of fast-starts in single and multiple hypoxic protocols

Latency distribution of control and experimental fast-starts for the both protocols are shown in Fig. 6A, B. In both protocols seventy-eight percent of all the latencies fall between 7.5-12.5 ms.

**Figure 6.**
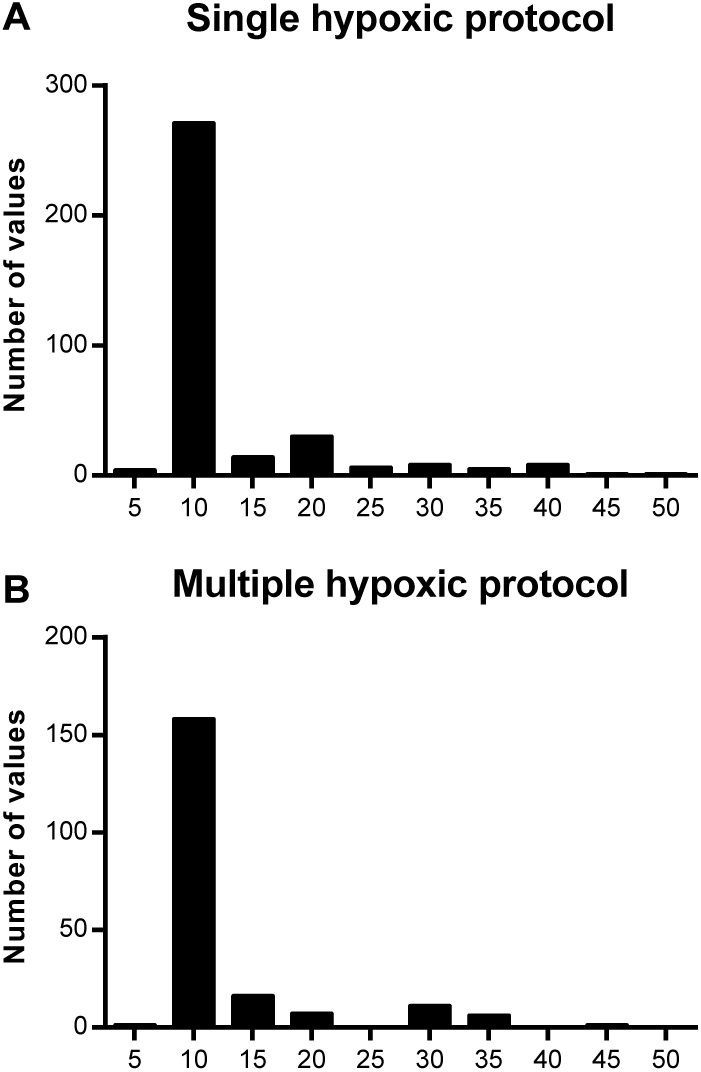
Distribution of startle response latencies. For both single and multiple hypoxia protocols, latency was measured as the first detectable movement of the head at the onset of a sound stimulus. **A) Latency distribution of control and experimental startle responses for the single hypoxic protocol.** Seventy-eight percent of the latencies fall between 7.5-12.5 ms. The average latency for this protocol was 12.4 ms ± 0.49 (mean ± SE) **B) Latency distribution of control and experimental startle responses for multiple hypoxic treatment.** Seventy-nine percent of the latencies fall between 7.5-12.5 ms. The average latency for this protocol was 12.7 ms ± 0.39 (mean ± SE)

### Morphological and electrophysiological identification of the M-cells

Mauthner cells were located about 300 µm below the surface of the medulla oblongata. The left and right cells from one fish are shown in Fig 7. The axons of these neurons are out of the plane of these 15 µm sections, and, as a result, we have placed a line to represent the trajectory of the axons. These large neurons have a composite axon cap with a central core and a peripheral portion surrounded by glia (only the glia nuclei are seen in these light micrographs). PHP processes can also be seen outside the glial layer (see Bierman et al., 2009).

**Figure 7.**
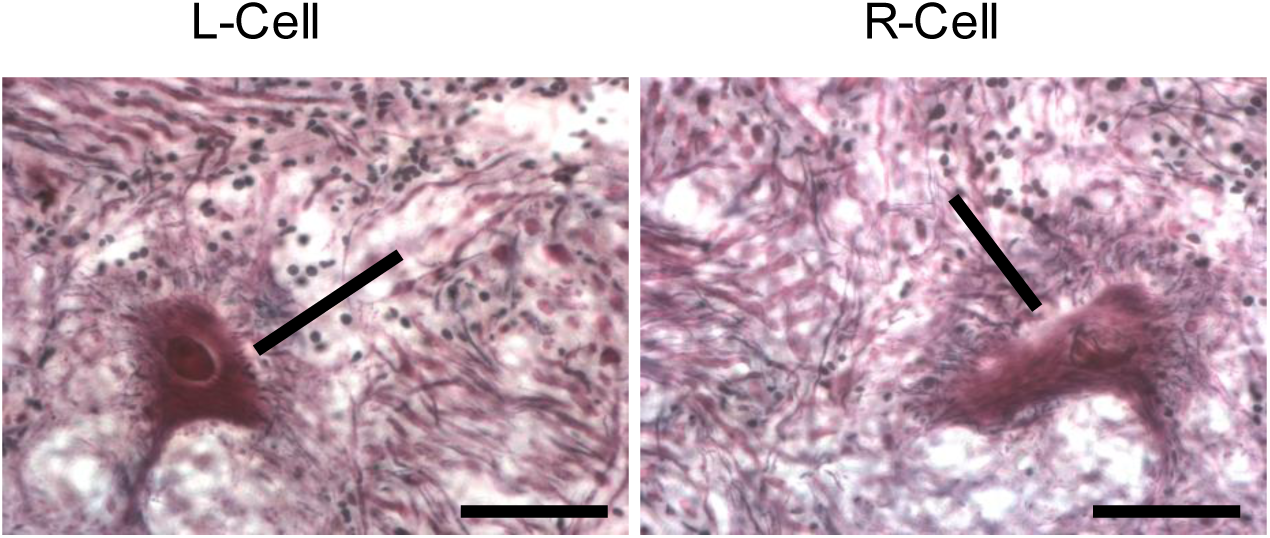
Morphological identification of the left and right Mauthner cell in a white grunt. Transverse sections (15 µm) at the level of the M-cells in the medulla oblongata. **A) Left M-cell.** A line has been placed to show the approximate trajectory of the M-axon which is out of the plane of the section. The line passes through a portion of the axon cap. **B) Right M-cell.** A line has been placed to show the approximate trajectory of the M-axon which is out of the plane of the section. The line passes through a portion of the axon cap.

A vertical depth profile of the M-cell extracellular negative field potential (blue line) and extrinsic hyperpolarizing potential (red line; EHP) are shown in Fig 8A. The microelectrode was inserted from the surface of the medulla oblongata ventrally to a depth of 325 µm. The electrode was then withdrawn dorsally moving in 25 μm steps. Representative recordings from three of the sites are shown as inserts. The location of the largest extracellular negative spike and positive EHP is around 150μm ventral from the surface of the medulla oblongata. Maximum recordings of the extracellular negative spike and the EHP from the left and right cells of the same fish are shown in Fig 8B. Lowering the stimulation voltage below threshold highlights the all-or-none nature of these potentials (stimulation rate, 1/s). Increasing the stimulus frequency from 1/s (upper trace) to 4/s (middle trace and lower trace) does not affect the all-or-none negative spike but does eventually eliminate the EHP (Fig. 8C). The M-cell extracellular negative field potential in the upper trace becomes larger when the electrode penetrates a neuron in the vicinity of the axon cap as seen in the lower trace (Fig. 8D). When one subtracts the field potential recorded intracellularly from that recorded extracellularly, there is a net negativity that is the so-called passive hyperpolarizing potential (PHP) that defines a PHP neuron.

**Figure 8.**
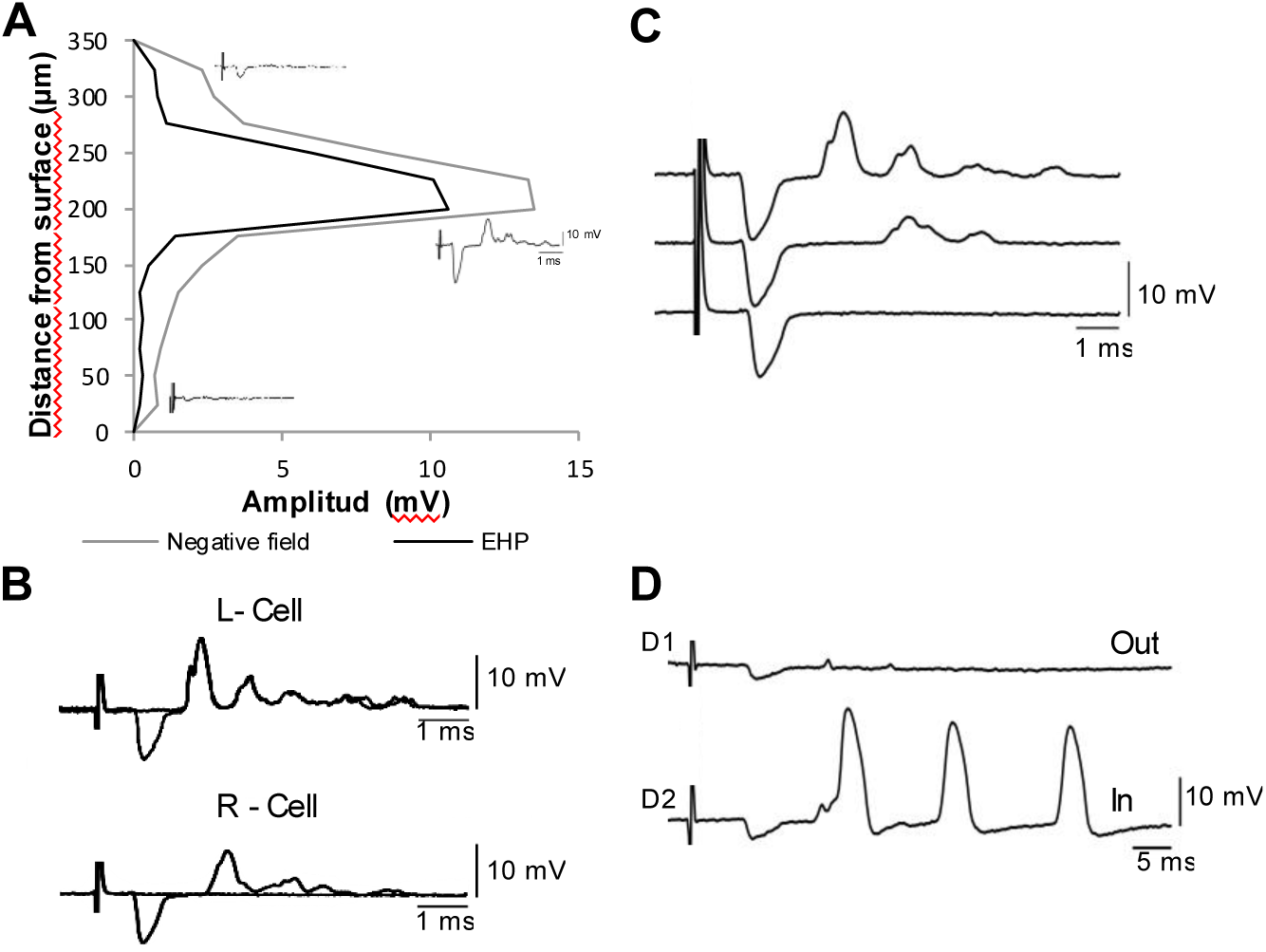
Electrophysiological characterization of the Mauthner cell of the white grunt. **A) Vertical depth profile of the M-cell extracellular negative field potential (grey line) and extrinsic hyperpolarizing potential (black line; EHP).** The electrode was inserted from the surface of the medulla oblongata ventrally to a depth of 325 µm. The electrode was then withdrawn dorsally moving in 25μm steps. Representative recordings from three of the sites are shown as inserts. The location of the largest extracellular negative spike and positive EHP is around 150μm from the surface of the medulla oblongata. **B) Maximum recordings of the extracellular negative spike and the EHP from the left and right cells of the same fish.** Lowering the stimulation voltage below threshold highlights the all-or-none nature of these potentials. Stimulation rate, 1/s. **C) Frequency Sensitivity of the EHP.** Increasing the stimulus frequency from 1/s (upper trace) to 4/s (middle trace and lower trace) does not affect the all-or-none negative spike but does eventually eliminate the EHP. **D) The passive hyperpolarizing potential neurons of the white grunt.** M-cell extracellular negative field potential in the upper trace becomes larger when the electrode penetrates the cell as seen in the lower trace. When one subtracts the field potential recorded intracellularly from that recorded extracellularly, there is a net negativity that is the so-called passive hyperpolarizing potential (PHP) that defines a PHP neuron.

## DISCUSSION

Anthropogenic activities and increased water temperatures associated with climate change have contributed to an increase in hypoxic conditions in nearshore ecosystems worldwide (Diaz, 2001; Jackson, 2008; Kennish, 2002; Zhang et al. 2010; Kroon et al. 2012; Rabalais et al., 2009). An increase in occurrence of hypoxia has been reported throughout the Caribbean where more than 25 eutrophic and hypoxic coastal zones have been identified (Diaz, et al., 2011; Ellison and Farnsworth, 1996). Condado lagoon water quality data indicates that values between 60-80% DO have become more common and that 40% DO (2.5 mg L^-1^) is the lowest recorded hypoxic event to date. Although no dissolved oxygen levels below 40% DO have been reported, a pattern of increasing frequency of low dissolved oxygen events has been documented in the past few years, mainly during Puerto Rico’s wet season (Lugo et al, 2011). An increase in hypoxic events has important management and conservation implications not only for the Condado lagoon but also the other four lagoons in the San Juan Bay Estuary system with poorer water quality.

Here we show for the first time that exposure of the white grunt (*Haemulon plumieri*), a tropical fish, to hypoxia significantly reduces the frequency of fast-starts an effect that lasts when a fish is returned to normoxic conditions. Since fast-starts are thought to be important for escape from predation, the survival of the white grunt and possibly other organisms in Condado lagoon is compromised with the potential for the disruption of population structure and dynamics.

A single exposure of white grunts to oxygen levels of 2.5 mg L^-1^ (40% DO) resulted in a decrease in frequency of fast-start responses and, the effect lasted for 24 h after exposure to low oxygen levels. The lack of a control for the 24 h exposure period does not allow us to eliminate habituation or handling as factors, but both are unlikely to have contributed to the observed results as we did not see those effects in controls during the treatments. The multiple hypoxic protocol was used to simulate the varied oxygen concentrations that white grunts might routinely encounter in the Condado lagoon. The frequency of fast-starts was significantly reduced when fish were tested 1 h after being transferred from hypoxic to normoxic conditions. We speculate that the lowest oxygen level of 3.7 mg L^-1^ (60% DO) used in the multiple hypoxic protocol is primarily responsible for changes in fast-starts although we cannot eliminate cumulative effects. The continued effect of hypoxia once a fish is returned to normoxic conditions is surprising and has far-reaching implications for fish survival even when exposed to mildly hypoxic conditions for short periods of time.

Differences in tolerance to hypoxia among fish are well known (Richards, 2011). A dissolved oxygen (DO) level of 2 mg L^-1^ has been used to define hypoxia that can result in the impairment of fisheries (Diaz, 2001; Vaquer-Sunyer and Duarte, 2008), however Vaquet-Sunyer and Duarte (2008) point out that this level underestimates sensitivity thresholds for most benthic organisms and that.4.6 mg L^-1^ would be more representative. Temperate fishes studied to date showed decreased startle responsivity at DO levels below 1.5-1.9 mg L^-1^ (golden grey mullet, *Liza aurata*, Lefrançois et al., 2005; European sea bass, *Dicentrarchus labrax*, Lefrançois and Domenici, 2006). We report similar behavioral effects but at higher oxygen concentrations than those reported for temperate fish. A possible explanation for the greater sensitivity of the white grunt to hypoxia may relate to higher water temperatures associated with tropical environments and the resultant decrease in oxygen availability. However, Rogers et al. (2016) have shown that tropical fish have a lower critical oxygen level (the oxygen level below which an organism cannot survive) than temperate species and are thus more tolerant of hypoxia. Other factors such as anthropogenic contaminants in Condado lagoon may contribute to the sensitivity of the white grunt to hypoxia. The response of white grunts from well-oxygenated, uncontaminated water to hypoxia will aid in the understanding of how concurrent stresses impact sensitivity.

In addition to physiological adaptations, fish have evolved behavioral adaptations to low oxygen environments that can increase fish tolerance to oxygen stress (Chapman and McKenzie, 2009; Ekan, 2010; Mandic et al., 2008; Richards, 2009, 2011; Wells, 2009). Many fishes use aquatic emergence (air-breathing) or aquatic surface respiration (ASR) as a strategy to counteract hypoxia (reviewed in Chapman and McKenzie, 2009; Kramer and Mehegan, 1981; Kramer, 1987; Kramer 1987; Lewis, 1970; Shingles, et al., 2005). Ninety-four percent of tropical freshwater fish studied utilized ASR under hypoxic conditions (Kramer and McClure, 1981) and 72% of species from marine habitats subject to hypoxia used this strategy (Kramer, 1983). Branchial respiration near the water surface increases the ability of fish to extract oxygen and creates a variable that can confound the relationship between hypoxia and behavioral changes. We did not observe ASR or aerial emergence by the white grunt during any phase of the single and multiple hypoxic protocols. As a result, hypoxia levels in this study were not altered by extraction of oxygen from the water surface or air.

A decrease in the ability to extract oxygen puts stresses on all organ systems, some of which may impact startle response behavior. Some examples of behavioral effects of hypoxia include decreased locomotor activity (Aboagye and Allen, 2014; Cannas et al., 2012), reduced feeding (Gamperl and Driedzic, 2009; Stierhoff, et al., 2006; Chabot and Claireaux, 2008), changes in dominance hierarchy (Sneddon and Yerbury, 2004) and reduced schooling behavior (Domenici, et al., 2002; Lefrançois, et al, 2009). Some physiological effects of hypoxia include changes in cardiovascular function (Shingles, et al., 2005), in respiratory patterns (Cannas, et al., 2012; Perry et al., 2009; Saint-Paul, 1984; Wannamaker and Rice, 2000), in reproduction and development (Wu, 2009) and in digestion (Wang et al., 2009). Other effects of hypoxia are related to oxygen uptake and include changes in gill structure (reviewed in Harper and Wolf, 2009) hemoglobin binding affinities (Wells, 2009) and tissue oxygen demands (Chabot and Claireaux, 2008; Hopkins and Powell, 2001). The short hypoxic exposure times used in this study would most likely affect respiration and cardiovascular function and possibly locomotor activity. Whether these possible changes could affect frequency of fast-starts is doubtful, although we cannot eliminate them as factors at this time.

We chose to define fast-starts as those occurring with latencies of 50 ms or less from stimulus onset to first movement. The average for all control and experimental latencies for the single hypoxic protocol was 12.65 ±7.36 ms (mean ± SE) and the average for the multiple hypoxic protocol was 12.41 ± 6.92 ms. These latencies are similar to auditory-evoked fast-start latencies of goldfish (12.40 ± 0.50 ms, Mirjany et al., 2011). Fast-starts were evoked by an abrupt sound stimulus (1 cycle of 100 Hz), a stimulus similar to that known to activate M-cells in goldfish (2 cycles of 200 Hz, Zottoli, 1977). It is unlikely that stimulation of the lateral line activated the M-cells. Although the lateral line innervates the goldfish M-cell with both excitatory and inhibitory components (Faber and Korn, 1975; Korn and Faber, 1975; Mirjany and Faber, 2011), inactivation of lateral line hair cells with CoCl2 or gentamicin does not change the probability of eliciting fast-start responses (Mirjany et al., 2011). Based on the short latency of responses, we speculate that many if not most of the responses to the sound stimulus are M-cell initiated and that the M-cells are activated by way of saccular afferents from the ear. We cannot exclude that some of the longer latency responses might be non-M-cell initiated (Liu and Fetcho, 1999; Zottoli et al., 1999) and involve the two pairs of M-cell homologs found caudal to the M-cell in segments five and six (Kohashi and Oda, 2008; Lee, et al., 1993; Nakayama and Oda, 2004). These factors, however, do not affect the conclusions of the present study. We did not see any impacts of the hypoxic treatments on the latency of fast-starts, which is consistent with results from previous studies (Lefrançois et al., 2005; Lefrançois and Domenici, 2006). Fast-starts elicited by an abrupt acoustic stimulus are initiated by M-cells and associated neurons (Eaton, et al., 1977, Zottoli, 1977). This implies that once the M-cell is brought to threshold, the timing of the remaining circuitry responsible for fast-starts (i.e., from M-cell to muscle) is not significantly affected by hypoxia. Although M-cells receive input from many sensory systems, the most powerful one is from large afferents that receive input from saccular hair cells (Furukawa, 1978; Lin et al., 1983; Zottoli et al., 1995). Limiting oxygen circulating over the gills results in a reduction of the sound-evoked, excitatory postsynaptic potential at the synapse between saccular hair cells and afferent fibers. Presynaptic mechanisms within hair cells appear to underlie this reduction (Suzue et al., 1987). If afferents are less responsive to sound stimulation, the probability that the M-cell will reach threshold is lessened and could explain the reduced frequency of fast-starts on exposure to hypoxia. This speculation will require further investigation. We have morphologically identified white grunt M-cells as a preliminary step to determine the site(s) affected by hypoxia in the fast-start circuit. The presence of a composite axon cap suggested that we would be able to find the cells electrophysiologically by the signature antidromically activated negative field potential followed by an extrinsic hyperpolarizing potential (Bierman et al., 2009; Zottoli et al., 2011). Indeed, such potentials were recorded along with evidence for the presence of passive hyperpolarizing potential (PHP) neurons. Future experiments will allow localization of the site or sites affected by hypoxia. Studying the M-cell and its circuit under hypoxic conditions will add insight into how low oxygen levels impact a single cell, the behavior it initiates, and ultimately how changes in the circuit might affect population and ecosystem levels.

Studies have shown that hypoxia can have a negative impact on species richness and abundance (Killgore and Hoover, 2001). Species that inhabit ecosystems like Condado lagoon at early life stages (e.g., eggs and larvae) will be susceptible to oxygen stress since they have limited mobility and thus can’t easily escape hypoxic conditions (Levin et al., 2009). Many adult and juvenile fishes, however, are able to detect and avoid hypoxic conditions (Jones, 1952; Karim et al., 2003; Wannamaker and Rice, 2000) with resultant changes in distribution (Pihl et al., 1991). Although staying in a hypoxic environment can convey an advantage to a predator of DO-stressed prey (Diaz and Breitburg, 2009), more often fish move to avoid hypoxia despite the increased risk of predation due to the loss of protective cover (reviewed in Chapman and McKenzie, 2009; Wolf and Kramer, 1987). Since the effects of low DO last beyond the hypoxic exposure, fish that move to normoxic conditions are subject to increased predation. In this study we examined a single, sub-lethal stressor, but multiple stressors may be acting at the same time (e.g., decreased pH, increased temperature and exposure to toxic pollutants; Somero et al., 2016). We may therefore be underestimating the possible impacts of environmental changes on the responsiveness and survival of fishes, and thus the more far-reaching effects on the distribution, abundance and diversity of fish and other species in complex nearshore marine habitats.

## List of Symbols and Abbreviations

ASR-: Aquatic surface respiration
DO: Dissolved Oxygen
M-cell-: Mauthner Cell
NEP -: Environmental Protection Agency National Estuary Program
SJBE-: San Juan Bay Estuary
SJBEP-: San Juan Bay Estuary Program

## ACKNOWLEDGMENTS

We thank Steve Treistman and the Institute of Neurobiology in Old San Juan, Puerto Rico for hosting S.J.Z. and Dr. Hector Marrero for his support in the laboratory. We also thank undergraduate students at UPR for their help in the collection, care of fish and in analyzing videos.

### Competing interests

The authors declare no competing or financial interests.

### Author contributions

The experiments were designed by S.J.Z., M.S.G. and L.R.M. M.S.G. and S.J.Z. performed the experiments. The paper was drafted, reviewed, and revised by M.S.G, L.R.M and S.J.Z. All authors commented on the manuscript and approved the submitted version.

### Funding

Funding This research was supported by the Puerto Rico Center for Environmental Neuroscience, National Science Foundation Centers of Research Excellence in Science and Technology Grant (HRD-1137725)

## Tables

**Table 1.** Timetable of single hypoxic treatment protocol

| **Treatments** | **Time** |
| --- | --- |
| *Tank acclimation* | *30 min* |
| *Baseline normoxia* | *18 min* |
| *Change in dissolved oxygen* | *25 min* |
| *Hypoxia control/ Hypoxic treatment* | *18 min* |
| *Change in dissolved oxygen* | *25 min* |
| *Reversal normoxia* | *18 min* |
| *Holding tank* | *24 h* |
| *Tank acclimation* | *30 min* |
| *24 h normoxia* | *18 min* |
| **Total time in chamber** | **3 h** |

**Table 2.** Timetable of multiple hypoxic treatment protocol

| **Treatments** | **Time** |
| --- | --- |
| *Tank acclimation* | *30 min* |
| *Baseline normoxia* | *30 min* |
| *Change in dissolved oxygen* | *25 min* |
| *Hypoxia control 1/ Hypoxic treatment 1* | *30 min* |
| *Normoxia* | *25 min* |
| *Change in dissolved oxygen* | *25 min* |
| *Hypoxia control 2/ Hypoxic treatment 2* | *30 min* |
| *Normoxia* | *25 min* |
| *Change in dissolved oxygen* | *25 min* |
| *Hypoxia control 3/ Hypoxic treatment 3* | *30 min* |
| *Normoxia* | *25 min* |
| *Reversal normoxia* | *30 min* |
| **Total time** | **5.55 h** |

